# Unusual photochemical characteristics of a novel BLUF-like protein from fungus

**DOI:** 10.64898/2026.08.14.744829

**Authors:** Saurabh Tewari, Suneel Kateriya

## Abstract

Blue light using Flavin (BLUF) proteins are microbial photoreceptors that are involved in various physiological responses. Their occurrence and biochemical properties in fungi remain poorly understood. Here, we investigated a putative BLUF photoreceptor from the corn-smut fungus *Mycosarcoma maydis* (MmBLUF). Domain analysis, multiple sequence alignment of BLUF core regions, and structural modelling indicated conserved canonical BLUF fold and flavin-pocket residues. However, when heterologously expressed, UV-visible and fluorescence spectroscopy revealed different spectral behaviour than canonical BLUF protein. Further, we tested the role of extended N-terminus in modulation of chromophore binding by expressing N-terminus truncated protein variants. Our results suggest that the unusual spectral behaviour is not linked to the truncation construct (extended N-terminus), which also showed similar spectral features, indicating that the extended N-terminus is unlikely to account for an unusual photodynamics characteristics. Our findings support MmBLUF as a structurally conserved putative fungal BLUF-like photoreceptor with different photochemical properties. Further studies are required to establish its chromophore identity, photocycle and function of this unusual BLUF-like domain from fungal system.

## 1. Introduction

Light sensing is vital beyond photosynthesis, and therefore organisms have developed various photoreceptors to sense the complete spectrum of light in order to gain unique cues from their environment. Among them BLUF is the blue light sensing photoreceptor. The *Euglena gracilis* (PAC enzyme, cAMP production) and *Rhodobacter sphaeroides* (AppA, transcriptional control) (Masuda & Bauer, 2002; Iseki et al., 2002). BLUF domains are nearly 100 amino acids long and sense blue light using a flavin chromophore(Gomelsky & Klug, 2002). They are found in nearly 10% of prokaryotes and unicellular eukaryotes(Losi & Gärtner, 2008). They are categorized based on the presence or absence of effector domain. Group I – includes BLUF proteins with effector domains (eg. AppA, BlrP1) and Group II are small, mostly just BLUF domain (200 amino acids or less).(Park & Tame, 2017a). BLUF are unique from other photoreceptors in having proton-coupled electron transfer (PCET). BLUF domains have Tyr, Gln, and Met moieties conserved around the flavin chromophore (Park & Tame, 2017a). Upon blue light electron + proton transfer from Tyr to flavin, generating radical pair forms that recombine into the signaling state but with marked change in the H-bonding pattern. (Möglich et al., 2010b). They have characteristic ~10 nm red-shift in flavin absorption peak after illumination (Masuda & Bauer, 2002). Mutation in conserved residues may either block the photocycle or cause some delay in signaling state (Masuda et al., 2008). BLUF domains form oligomers, most commonly dimers, and exhibit a ferredoxin-like core structure. This fold consists of a five-stranded mixed β-sheet with both parallel and antiparallel arrangements in the order 4-1-3-2-5, accompanied by two α-helices that run parallel to the β-strands(Anderson et al., 2005).

A tryptophan residue present in the 5^th^ beta strand is partially conserved and is implicated in different proteins to be responsible for light induced confirmational change and further signaling by switching its position with a conserved methionine residue after photoexcitation (Jung et al., 2006). C-terminal caps have also been found to be involved in signal transduction in BLUF proteins (Grinstead et al., 2006), and its role in overall stability has also been tested. (Tanwar et al., 2016).

Fungi play a vital role in the recycling of organic matter, act as significant pathogens in both plants and animals, and are widely exploited in industry for producing secondary metabolites and catalytic enzymes. Therefore, progress in fungal biology research is essential for deepening our knowledge of their pathogenic mechanisms and enhancing their biotechnological applications. It has been shown that fungal genome expression is intimately involved with light and it controls major life cycle events like sporulation, fruiting body formation, secondary metabolite production, phototropism etc.(Ruger-Herreros et al., 2011) (Chen et al., 2009). Even though BLUF photoreceptors have been described in many fungi, their mechanistic study is lacking. (Yu & Fischer, 2019). Recently, the fungal photoreceptor research has focused on WC and phytochromes, leaving BLUF diversity underexplored in fungi (Bayram & Bayram, 2023). When fungi are exposed to blue light, their transcriptome can respond on a massive scale — sometimes with tens of percent of the entire genome becoming active, as seen in edible mushrooms. This suggests that fungi may have additional blue-light sensors, possibly including BLUF proteins, which could be important if they link into signaling pathways (Zhu et al., 2024). We can therefore conclude that primary characterization of BLUF protein in fungi would be the first step towards understanding their structure, photocycle, and possible role they may play in Fungi. This could be important in pathogenic response understanding and other growth-related aspects, along with the possibility of optogenetic application. In the present study we identified a BLUF domain containing protein (MmBLUF) from *Mycosarcoma maydis* which causes Smut disease in corn. We studied photochemical properties of the protein and the effect of extended N-terminus on this protein.

## 2. Materials and methods

### 2.1 Sequence retrieval and bioinformatic analysis of BLUF domains

The putative MmBLUF sequence (protein accession XP_011386128.1) was identified using BLAST from NCBI (http://www.ncbi.nlm.nih.gov/). Conserved domain in the protein sequence was identified using the Conserved Domain Database in NCBI via CD search (Wang et al., 2023). Basic biophysical characteristics were predicted using the ProtParam Tool (Gasteiger et al., n.d.). Homology analysis was done using sequences retrieved from the NCBI server (O’Leary et al., 2024) employing ClustalW (Thompson et al., 1994), and results were visualized using Jalview (A. M. Waterhouse et al., 2009). Secondary structure was predicted using PSIPRED (McGuffin et al., 2000), JPRED4 (Drozdetskiy et al., 2015), and NetSurf (Høie et al., 2022). The predicted structure was incorporated into an image using BioRender (BioRender.com).

### 2.2 Structural modelling and comparative structural analysis of MmBLUF

The structure of the MmBLUF core was modelled using SWISS-MODEL. An independent structural prediction was also obtained using the AlphaFold server for comparison of the overall fold. For comparative structural analysis, the SWISS-MODEL-generated MmBLUF core was superimposed with the crystal structure of the AppA BLUF domain from *Rhodobacter sphaeroides* (PDB ID: 2IYG, chain A) using the CEalign algorithm in PyMOL. Alignment coverage, root mean square deviation (RMSD), and number of aligned Cα atoms were used to understand structural similarity between the canonical and fungal protein. The FMN cofactor coordinates from holo-AppA were used to examine the geometric proximity of the predicted MmBLUF flavin-binding pocket. MmBLUF residues located within 3.5 Å of the transferred FMN were identified using PyMOL.

### 2.3 Phylogenetic Analysis of BLUF proteins

BLASTp programme of NCBI was used to find putative fungal BLUF candidates using the MmBLUF core domain region, and canonical BLUFs were selected based on the literature. Only the predicted BLUF-core regions were used for analysis. The amino acid sequences were aligned using MUSCLE in MEGA software. Using the Find Best DNA/Protein Models (ML) function in MEGA, the best amino acid substitution model was the LG+G+I model due to its lowest Bayesian Information Criterion score. A maximum-likelihood phylogenetic tree was constructed under the LG+G+I model, and branch support was assessed using 1000 bootstrap replicates. The final tree was also visualised and edited in MEGA.

### 2.4 Cloning, heterologous expression, and purification of MmBLUF

MmBLUF gene was subcloned into the pASK-IBA43 plasmid from the pUC57 plasmid. pASK-MmBLUF was transformed into BL21(DE3). Transformed cells were grown in Terrific Broth (TB) and induced with 200 μg/L anhydrotetracycline (IBA, Göttingen, Germany). Cells were harvested by centrifugation, resuspended in lysis buffer, and then sonicated to form a cell lysate, which was then centrifuged to get soluble fraction. The soluble fraction was allowed to bind to Ni-NTA beads, and His-MmBLUF was later eluted using 200 μM imidazole.

### 2.5 Site-directed mutagenesis and N-terminal truncation of MmBLUF

For N-terminal truncation, genes named MmBLUF (60-294), MmBLUF (100-294) were PCR-amplified using the MmBLUF plasmid as a template with primers listed in Table S1 and cloned into the pASK-IBA43 plasmid. For site-directed mutagenesis, T93Y mutation primers were used to amplify the entire plasmid using Phusion polymerase (Thermofischer Scientific), followed by DpnI digestion of the parental DNA. All constructs were confirmed by automated DNA sequencing.

### 2.6. UV-Vis spectroscopic analysis of protein

For UV-Visible spectroscopy, the eluted protein was purified by exchanging the imidazole buffer for PBS (100 mM) via dialysis. UV-visible spectroscopy was performed using an Agilent 3000 UV-VIS spectrophotometer. UV-Visible spectroscopy of the protein was done in the 300-600 nm wavelength range. The data obtained were plotted using GraphPadPrism software.

### 2.7 Fluorescence Spectroscopy of purified MmBLUF and N-terminal truncated MmBLUF

Steady state fluorescence spectroscopy was carried out using a Perkin-Elmer fluorescence spectrophotometer. The protein sample was excited at 390 nm. Fluorescence emission was recorded between wavelengths of 400nm to 650 nm in the dark. The data obtained were plotted using GraphPadPrism software.

### 2.8 Immunoblotting of purified recombinant MmBLUF

Immunoblotting was performed to detect His-tagged protein expression. Protein samples were first resolved by SDS-PAGE and then transferred onto a nitrocellulose membrane. The membrane was blocked with 5% fat-free milk to prevent non-specific antibody binding, followed by incubation with a penta-His primary antibody (1:5000 dilution). After washing with 1× PBS containing 0.1% Tween-20, the membrane was incubated with an HRP-conjugated anti-mouse secondary antibody (1:5000 dilution). The protein signals were then detected using a chemiluminescence-based detection method.

## 3. Results and discussion

### 3.1 MmBLUF is a BLUF-like photoreceptor with an extended N-terminus

In the present study we found that MmBLUF is a BLUF-like photoreceptor with an extended N-terminus. The homology and structure prediction analysis revealed that MmBLUF have conserved canonical BLUF chromophore binding residue. Conserved-domain analysis of the MmBLUF(294 amino acid long) showed a central region from 115-206 amino acids having BLUF domain-like features, preceded by an N-terminus region of 115 amino acids and followed by a C-terminus domain of around 88 amino acids with no similarity predicted for any other established domains (Figure 1A). The MmBLUF core was aligned with representative canonical BLUF proteins, including AppA, Slr1964, YcgF, BlrB, Tll0078, OaPAC, and bPAC. The alignment showed distributed conservation across the approximately 93-residue core rather than conservation limited to a short local segment (Figure 1B). In the BLUF domain, signal translation is determined by the conserved tyrosine, glutamine, and tryptophan (or methionine) residues that interact directly with the flavin chromophore (Yuan et al., 2006). MmBLUF retained residues corresponding to the characteristic BLUF Tyr–Gln– Met network, including Tyr6, Gln49, and Met92. Together, these sequence-level observations support the assignment of MmBLUF as a putative BLUF-like protein with an unusually extended N-terminus.

**Figure 1:**
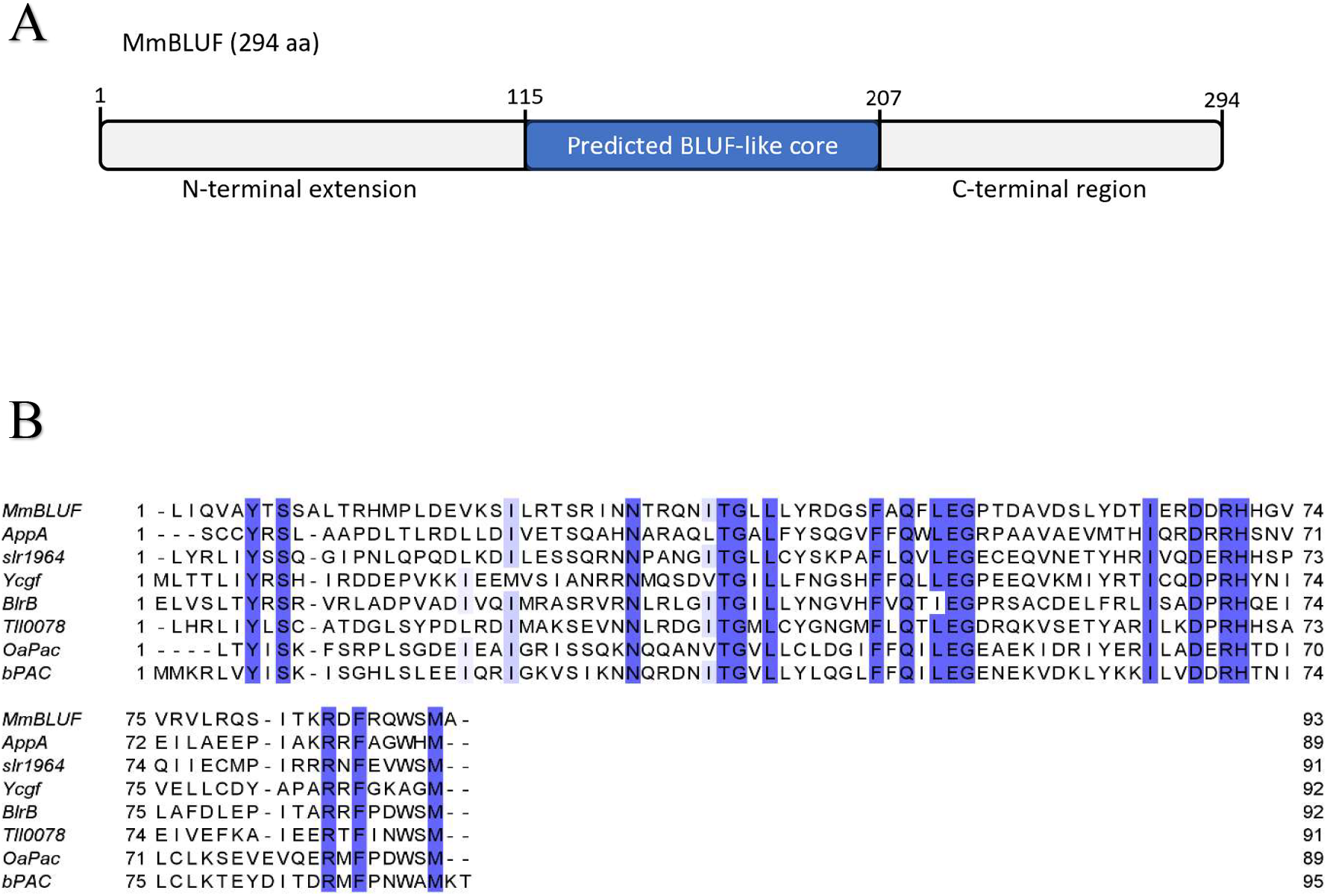
Domain architecture and sequence conservation of the predicted BLUF-like core in MmBLUF. (A) Schematic representation of the 294-amino-acid MmBLUF protein. (B) Multiple sequence alignment of the predicted MmBLUF BLUF-like core with representative canonical BLUF proteins. Conserved residues are shown in a blue background.

### 3.2 Phylogenetic placement of MmBLUF among fungal and canonical BLUF proteins

Phylogenetic analysis showed that MmBLUF grouped near BLUF-like proteins from closely related basidiomycete fungi, including *Moesziomyces aphidis* and *Moesziomyces antarcticus* (Figure 2). This placement is consistent with the taxonomic origin of MmBLUF and supports its classification as a putative fungal BLUF-like protein.

**Figure 2.**
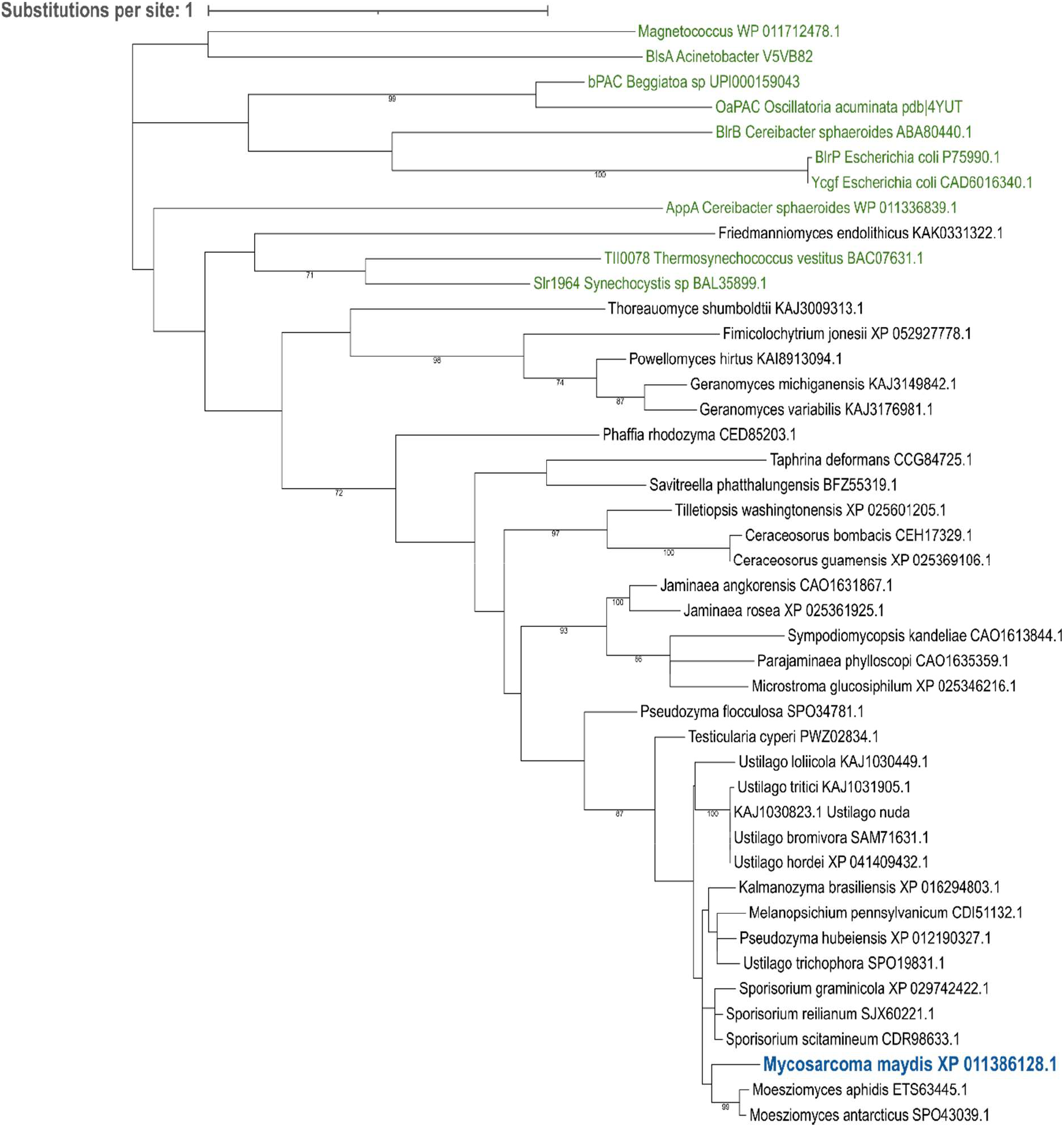
Maximum-likelihood phylogeny of MmBLUF, putative fungal BLUF-like proteins, and representative canonical BLUF proteins. Phylogenetic analysis was performed using the predicted BLUF-core regions of MmBLUF, selected fungal BLUF-like candidate proteins, and representative canonical BLUF proteins. The tree was inferred using the maximum-likelihood method under the LG+G+I amino-acid substitution model. Branch support was assessed using 1000 bootstrap replicates, and only bootstrap values ≥70% are shown. MmBLUF from Mycosarcoma maydis is highlighted in blue, while representative canonical BLUF proteins are shown in green. Putative fungal BLUF-like proteins are shown in black. The scale bar represents the estimated number of amino-acid substitutions per site.

### 3.3 Structural superimposition of Fungal and canonical BLUF AppA

Structural superposition of the predicted MmBLUF (Figure 3A) core with AppA produced an RMSD of 1.10 Å over 88 aligned Cα atoms, covering 94.6% of the MmBLUF model (Figure 3B). Visible overlap in both the alpha-helical and Beta-sheet region can be inferred as close structural similarity between canonical AppA and predicted BLUF. Furthermore, the AppA-derived FMN occupied a corresponding internal cavity in the MmBLUF model and was surrounded by putative pocket residues, including Tyr6, Gln49, and Met92 (Figure 3C). However, because FMN was transferred from the AppA holo structure rather than experimentally resolved or docked in MmBLUF, these observations demonstrate geometric compatibility of a putative flavin-binding pocket and do not confirm native FMN binding, cofactor occupancy, or the precise hydrogen-bonding arrangement (Figure 3D).

**Figure 3.**
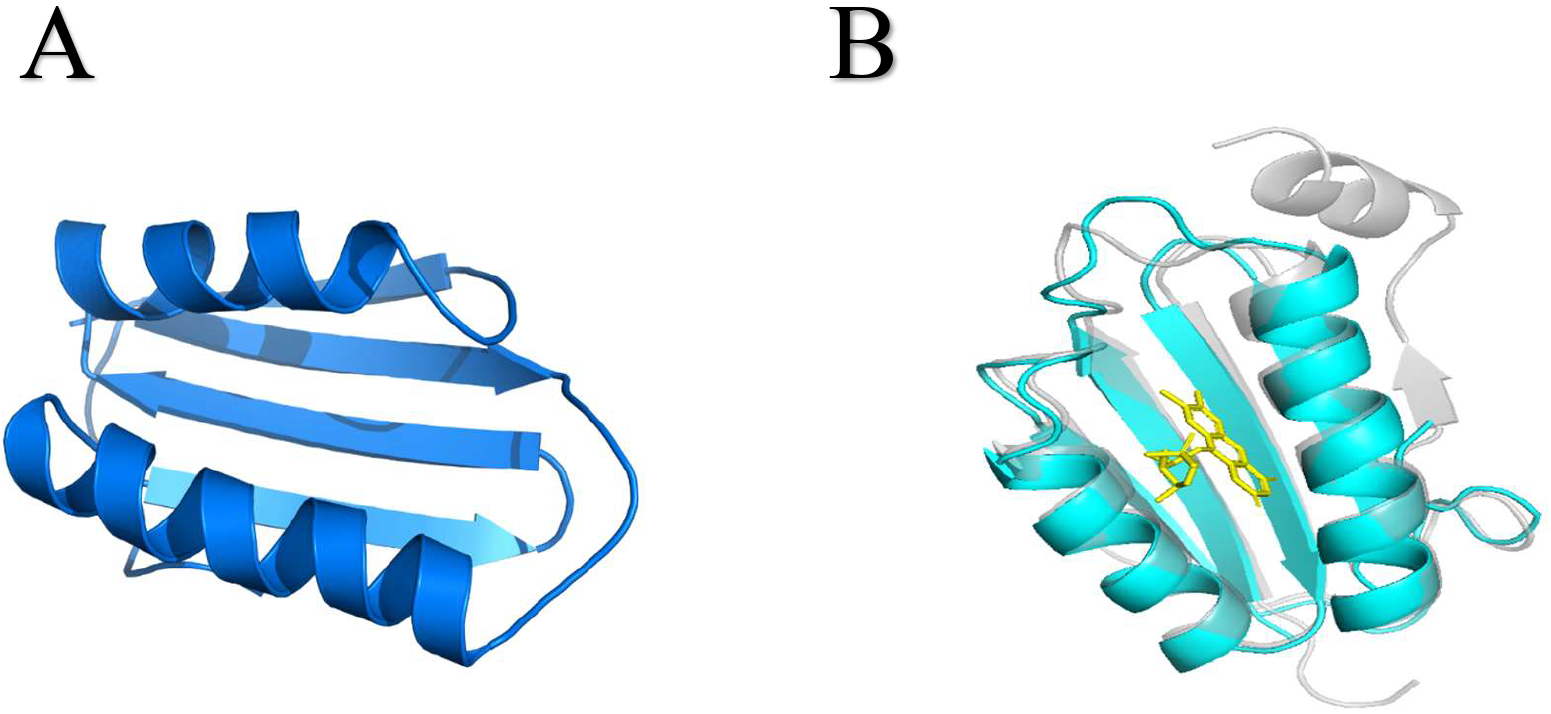
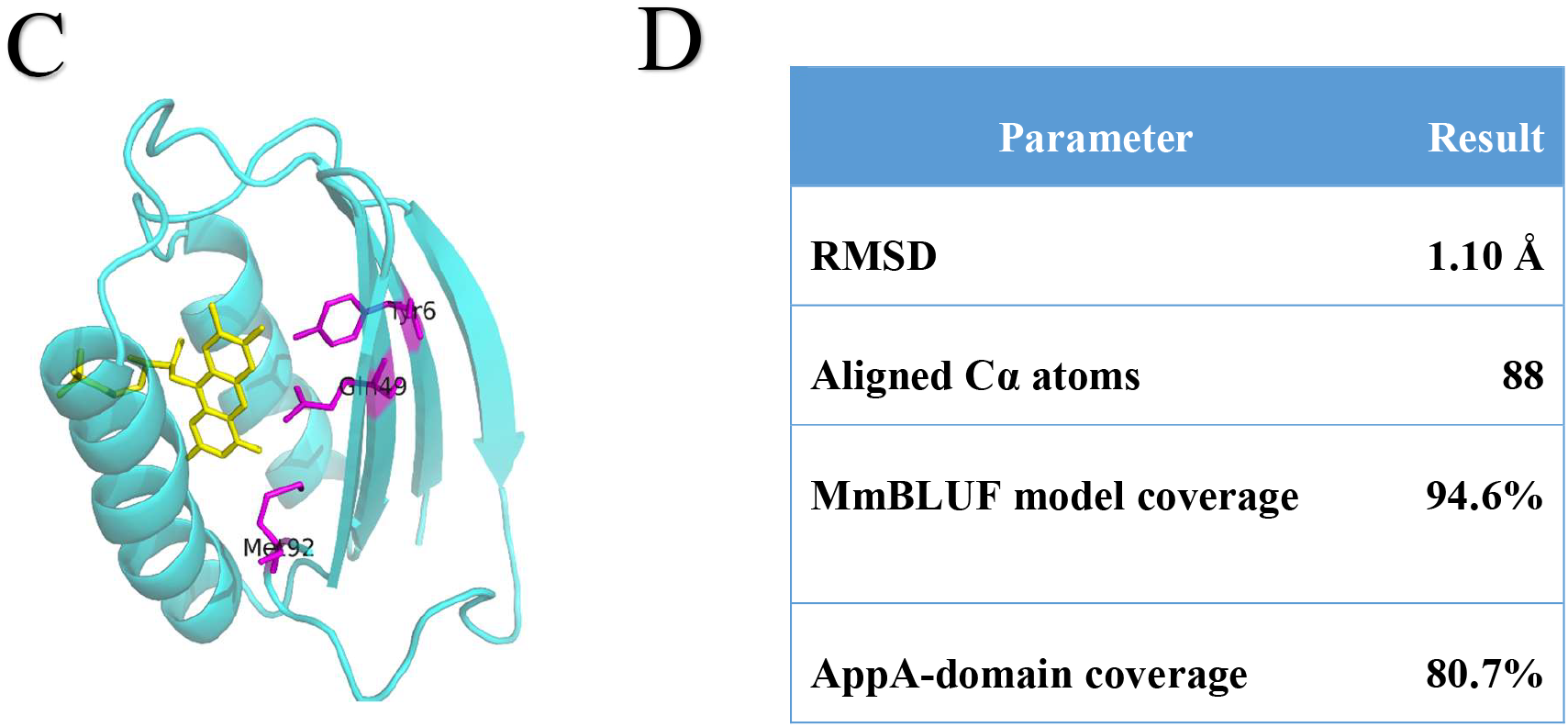
Structural comparison of the predicted MmBLUF core with the canonical AppA BLUF domain. **(A)** SWISS-MODEL-predicted structure of the MmBLUF BLUF-like core. **(B)** Structural superposition of the MmBLUF model (cyan) with the experimentally determined AppA BLUF domain from *Rhodobacter sphaeroides* (PDB ID: 2IYG, chain A; grey). The FMN cofactor retained from the AppA structure is shown as yellow sticks. **(C)** Close-up view of the putative flavin-compatible cavity in MmBLUF, showing the conserved Tyr6, Gln49 and Met92 residues as magenta sticks and the transferred AppA FMN in yellow. **(D)** Quantitative parameters obtained from the structural alignment, including RMSD, number of aligned Cα atoms and alignment coverage.

#### Spectral analysis of recombinant MmBLUF reveals non-canonical photodynamic features

In UV-Vis Spectroscopy, characteristic peaks of BLUF proteins are observed around 450nm and 370nm, for example, in AppA. Absorbance maxima are observed at 365nm and 445nm (Kraft et al., 2003). Wild-type MmBLUF, however, showed a broad, non-canonical absorption profile with a poorly resolved shoulder around 400–410 nm. No distinct maximum in the canonical BLUF absorption region near 440–450 nm was observed (Figure 4A). Across different BLUF proteins, the peaks vary: for Slr1964, the maximum absorbance is at 443nm, whereas for T110078 it is blue-shifted to 439nm (Okajima et al., 2005). Absorbance maxima could even be red-shifted, as in SnfB, where it is at 460nm (Tanwar et al., 2016b). On top of that, the absorbance characteristics of BLUF protein may even change due to mutations in the conserved residues around the Flavin-binding pocket. For example, AppA-Y21F mutant shows a blue shift in absorbance peak, from 365 to 359 and 445 to 444nm. Furthermore, a Y21L mutant showed even greater blue shift to 359 and 438nm. Another mutant Q63L shifts it to 437, but Q63E red shifts it to 449nm (Dragnea et al., 2010). The trend is that the more hydrophobic the binding pocket, the more blue-shifted the absorbance maxima are (Kraft et al., 2003). The normalized spectra of MmBLUF (60–294) and MmBLUF (100–294) (Figure 4B) remained broadly similar to that of the wild-type protein, indicating that removal of up to the first 100 N-terminal residues did not restore a typical BLUF-like spectral signature. The T93Y substitution was introduced at a non-conserved N-terminal position to test whether incorporation of an additional aromatic side chain influenced the spectral environment of the protein (Figure 4C). However, the normalized spectrum of the T93Y mutant largely overlapped with those of the wild-type and truncation constructs, with no substantial shift in the visible-region feature (Figure 4D). Figure S1 shows MmBLUF and free flavin spectra for comparison. These observations suggest that neither the extended N-terminal region nor the non-conserved residue at position 93 is the primary determinant of the atypical MmBLUF spectrum under the conditions tested. Another factor contributing to a non-canonical Absorption Spectrum could be the chromophore itself; for example, different oxidation states of flavin have different absorbance peaks. Free flavin in its oxidized form in H2O absorbs maximally at 445nm and 373nm. The same flavin in its reduced state absorbs maximally at 395 and 295nm, respectively (Ghisla et al., 1973). Similarly, FAD has also been shown to be bound to BLUF photoreceptors (Park & Tame, 2017b). FAD shows absorption peaks at 450nm and 375nm in its oxidized state, but the peaks quickly blue-shift to 410nm and 350nm in reduced state (Ghisla et al., 1973; Westphal et al., 2006). Figure S1 shows MmBLUF and free flavin spectra for comparison. However, the presence of MmBLUF was observed in the western blot images (Figure S2).

**Figure 4.**
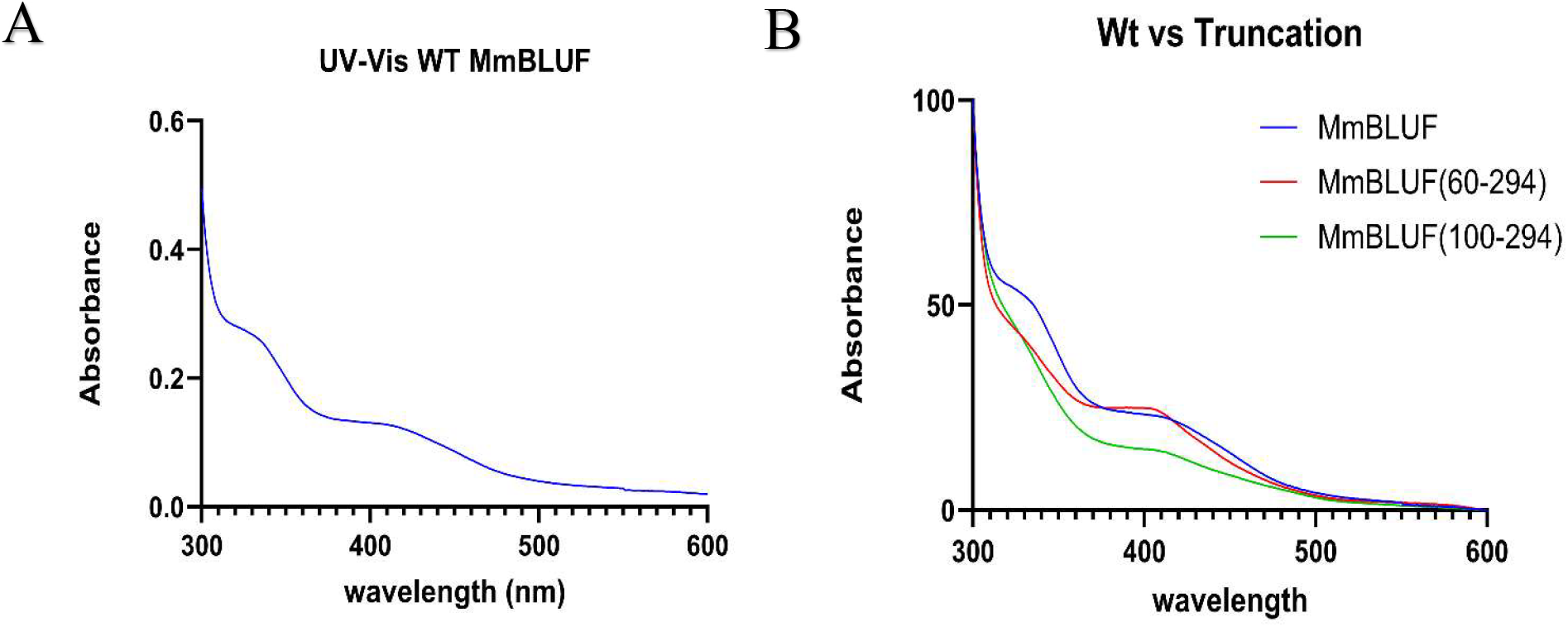
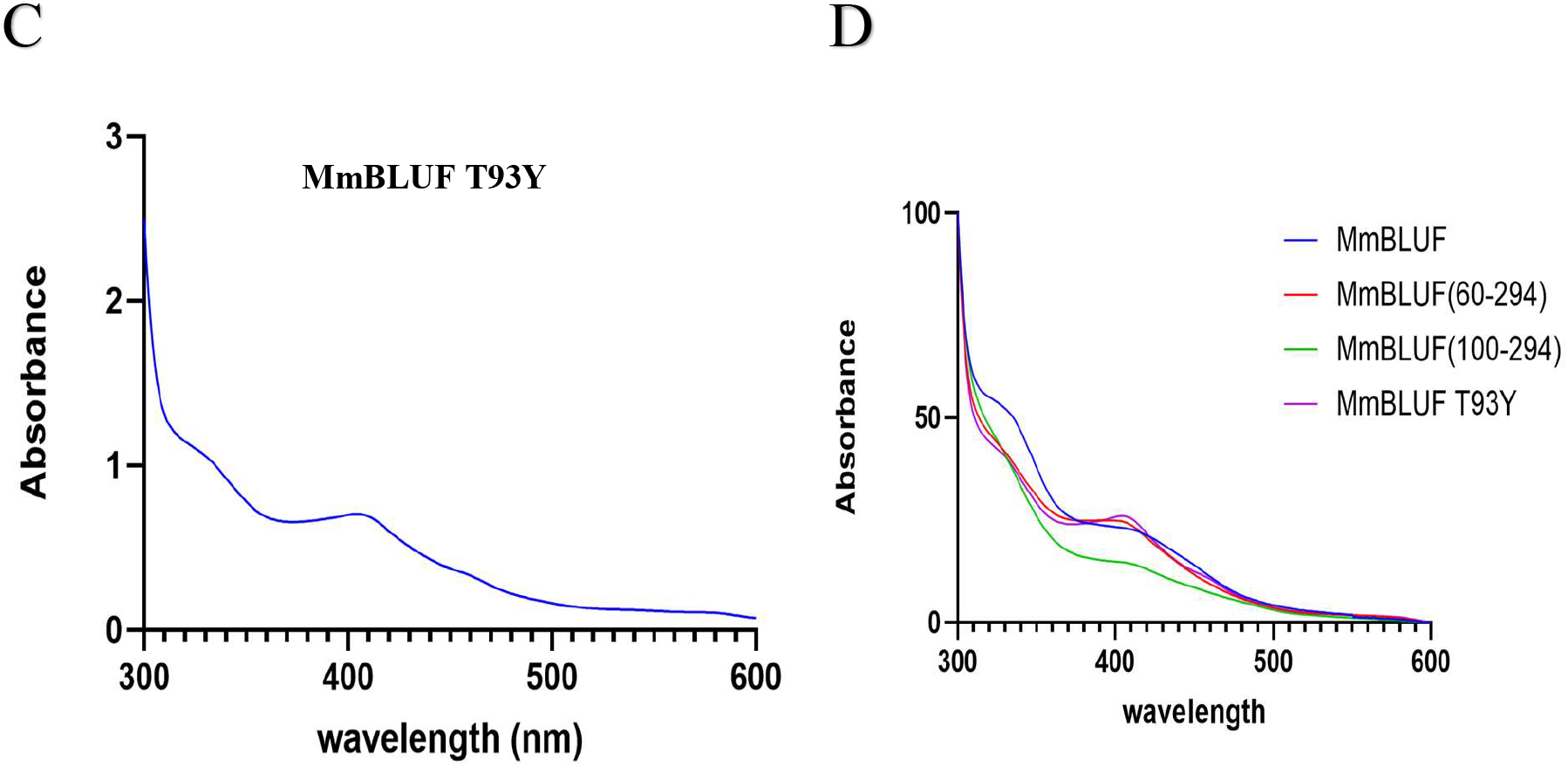
UV–visible absorption spectra of wild-type MmBLUF, N-terminal truncation variants, and the T93Y mutant of MmBLUF. **(A)** UV–visible absorption spectrum of wild-type MmBLUF recorded between 300 and 600 nm. **(B)** Normalized spectral comparison of wild-type MmBLUF with the N-terminal truncation variants MmBLUF (60–294) and MmBLUF (100–294). **(C)** UV–visible absorption spectrum of the MmBLUF T93Y mutant, in which a non-conserved threonine in the N-terminal region was replaced with the aromatic residue tyrosine. **(D)** Normalized comparison of wild-type MmBLUF, both N-terminal truncation variants, and the T93Y mutant.

In fluorescence spectroscopy, characteristic emission peaks are around 500nm. For AppA, it is 503nm (Dragnea et al., 2010). However, wild-type MmBLUF displayed a broad fluorescence emission band with a maximum in the approximately 450–470 nm region, which is markedly blue-shifted relative to the emission maxima generally reported for canonical flavin-bound BLUF proteins near 500 nm (Figure 5A). In emission spectra, the variation in peaks also arises due to various factors; for example, for AppA, it is 503nm (Dragnea et al., 2010), for Snfb, it is 516nm (Tanwar et al., 2016b). For the Q63L mutant of AppA, it red-shifts to 514 with a higher intensity, but for Q63E, intensity decreases, and the maximum absorbance is at 506nm. Similarly, the chromophore could also affect the fluorescence spectra; for example, the emission peak of oxidized FMN/FAD is around 520nm. However, a recent study concludes that reduced flavin is non-fluorescent.(McBride et al., 2023). The N-terminal truncation construct MmBLUF (60–294) and the T93Y mutant showed broadly similar emission profiles (Figure 5B, C). After normalization, the three spectra largely overlapped, with no substantial shift in the emission maximum or overall spectral shape (Figure 5D). These results indicate that removal of the first 59 residues and substitution of the non-conserved N-terminal Thr93 with an aromatic tyrosine do not appreciably alter the fluorescence signature of MmBLUF under the tested conditions. The atypical blue-shifted emission therefore appears unlikely to arise solely from the extended N-terminal region or from the local aromatic character at position 93.

**Figure 5.**
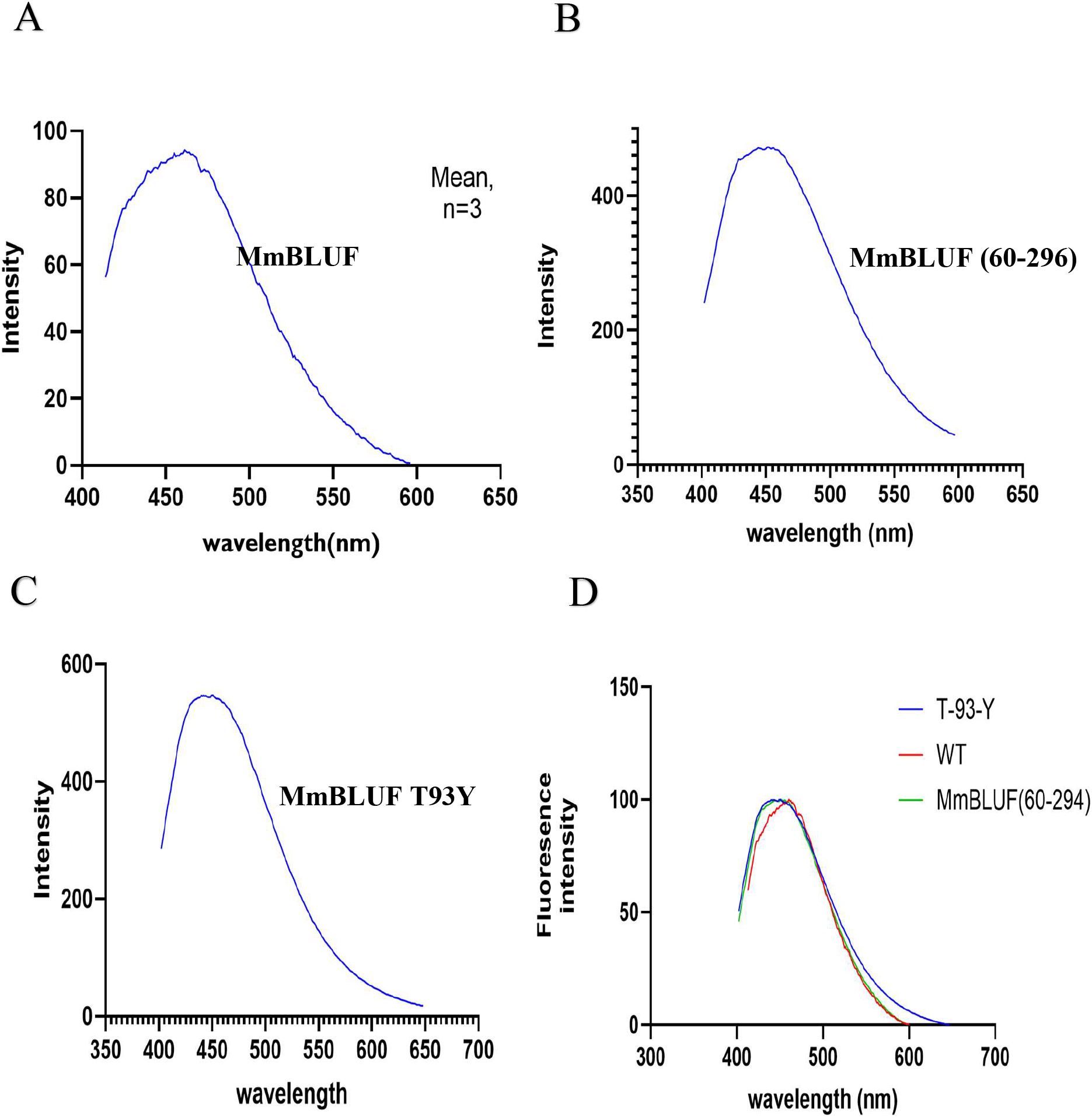
Fluorescence emission spectra of wild-type MmBLUF, an N-terminal truncation variant, and the T93Y mutant. (A) Fluorescence emission spectrum of wild-type MmBLUF. (B) Emission spectrum of the N-terminal truncation construct MmBLUF (60–294). (C) Emission spectrum of the MmBLUF T93Y mutant, in which a non-conserved N-terminal threonine was replaced with tyrosine. (D) Normalized comparison of the fluorescence emission spectra of wild-type MmBLUF, MmBLUF(60–294), and MmBLUF T93Y. Samples were excited at 390 nm, and emission was recorded between 400 and 650 nm.

## Conclusions

Overall our study concluded that the apparent blue-shift in wavelength maxima of absorbance and emission of MmBLUF cannot be attributed to a single factor, but rather different aspects have to be considered, including conserved residue analysis, chromophore characterization, and chromophore oxidation state. No significant change in the spectral properties of the mutant and wild type was observed. Even if there is a small difference, like those visible in the AppA mutants by shifts of a few nanometers (Dragnea et al., 2010). It would be hard to infer without consistent replicates. Therefore, further experiments are required to understand the photochemical properties and the mechanism behind the deviated behaviour.

## Author Contributions

S.K. conceived the project. S. T. designed the experiments in the guidance of S.K. and drafted the manuscript. All the experiments were performed by S. T. All authors reviewed and approved the final manuscript.

## Data Availability

Data will be made available on request.

## Acknowledgement

SK is grateful to STARS-MHRD (STARS/APR2019/BS/563/FS) and acknowledges the Department of Biotechnology for financial assistance through DBT-PG fellowship.

