## Supplementary materials for "Unusual photochemical characteristics of a novel BLUF-like protein from fungus"

**Protein and nucleotide sequence of the construct**

**Protein sequence**

MmBLUF (*Mycosarcoma maydis XP_011386128.1)*

MLQQASTSNPSLVPLRGQRPVVHLEHPPSVPRSASRGSFENGLATSLPPRRPPASFTRSASSHNNNIVTPAKRPISGHSSRQSSSDSIESVTTASSFTRASTSNYDSGSSDEEGLIQVAYTSSALTRHMPLDEVKSILRTSRINNTRQNITGLLLYRDGSFAQFLEGPTDAVDSLYDTIERDDRHHGVVRVLRQSITKRDFRQWSMAFRDLDMIKKCSSATSGSHHLGNEMTPEDAVNEGFS)LMNVGLRVGGPSDNEYRPDALGNAKLDLPADMSASMRRLVTTFYRFMDRPL

**Nucleotide sequence from NCBI**

>NC_026478.1:486892-487776 Mycosarcoma maydis chromosome 1, whole genome shotgun sequence

ATGCTACAACAAGCTAGCACCTCGAACCCTTCTCTTGTGCCTCTTAGAGGCCAGAGACCGGTTGTGCATCTGGAGCATCCACCTTCTGTCCCTCGTTCAGCTAGTCGCGGCTCTTTCGAAAATGGCCTTGCAACTTCTCTTCCACCAAGGCGGCCGCCTGCCTCCTTTACTCGCAGCGCTTCTTCGCACAACAACAACATCGTCACGCCTGCGAAACGACCAATTTCAGGTCACTCGTCCCGCCAGTCGTCGTCCGACTCGATCGAGTCAGTGACTACTGCCTCCTCCTTCACTCGCGCCTCAACTTCCAACTACGACTCTGGCAGCAGCGACGAGGAGGGGCTGATCCAGGTGGCCTACACCTCCTCAGCCCTAACCCGTCACATGCCGCTGGACGAGGTCAAGTCAATCTTACGTACTTCGCGCATCAACAATACCAGGCAAAACATTACTGGGCTACTTCTGTACCGTGATGGCTCATTCGCACAGTTTCTCGAAGGACCGACGGACGCTGTTGATTCGCTTTACGACACGATTGAGCGCGACGATCGTCACCACGGTGTTGTTCGCGTGCTTCGACAGTCGATCACTAAGCGTGATTTTCGTCAGTGGAGCATGGCGTTTCGCGATCTCGACATGATCAAGAAGTGCTCCTCTGCCACGTCCGGTTCCCACCACCTCGGTAATGAGATGACACCCGAGGACGCCGTCAATGAAGGCTTCAGTCAGCTCATGAACGTTGGTCTTCGTGTCGGTGGCCCTTCTGACAACGAGTATCGCCCGGACGCTCTCGGTAACGCCAAACTTGATCTACCAGCTGACATGTCTGCCAGTATGCGTAGACTCGTTACCACTTTCTACCGTTTCATGGACCGGCCCCTCTAA

**Synthesized construct- Nucleotide sequence**

ATGCTGCAACAAGCGTCTACTTCTAACCCGAGCCTGGGTCCTCTGCGTGGTCAGCGTCCTGTTGTACACCTGGAACATCCGCCGTCTGTACCGCGCTCTGCGTCCCGTGGCTCTTTTGAGAATGGTCTGGCCACCTCTCTGCCGCCGCGCCGTCCTCCGGCGAGCTTTACTCGTTCTGCGAGCAGCCACAACAACAATATCGTAACCCCAGCGAAACGTCCGATTTCTGGCCATTCCTCTCGTCAGTCTTCTTCTGATAGCATCGAATCCGTTACCACCGCATCCTCTTTCACTCGTGCATCCACCAGCAACTACGATTCTGGTTCTTCCGACGAAGAGGGTCTGATCCAGGTTGCTTACACGTCCAGCGCTCTGACTCGTCACATGCCGCTGGATGAAGTGAAGAGCATCCTGCGTACTTCCCGCATCAACAACACCCGTCAGAACATCACGGGCCTGCTGCTGTATCGTGACGGCTCCTTCGCACAGTTCCTGGAAGGTCCGACTGATGCCGTCGATTCCCTGTATGATACGATTGAGCGTGATGACCGTCATCACGGCGTCGTGCGTGTTCTGCGTCAGTCCATCACCAAACGTGACTTCCGTCAGTGGAGCATGGCTTTCCGTGATCTGGACATGATTAAAAAATGCTCTTCCGCTACCTCCGGCAGCCACCACCTGGGTAACGAAATGACCCCAGAAGACGCGGTGAACGAAGGCTTCTCCCAGCTGATGAACGTGGGTCTGCGCGTTGGTGGCCCGTCCGACAACGAATACCGCCCGGACGCACTGGGCAACGCCAAACTGGACCTGCCAGCAGACATGTCCGCTTCTATGCGCCGCCTGGTTACCACCTTCTACCGCTTTATGGATCGCCCGCTG

**Sequences of BLUF proteins used for sequence alignment and phylogenetic analysis**

**Core BLUF domain region has been extracted and present here, with accession number of full sequence**

>AppA_Cereibacter_sphaeroides_WP_011336839.1

SCCYRSLAAPDLTLRDLLDIVETSQAHNARAQLTGALFYSQGVFFQWLEGRPAAVAEVMTHIQRDRRHSNVEILAEEPIAKRRFAGWHM

>slr1964_Synechocystis_sp_BAL35899.1

LYRLIYSSQGIPNLQPQDLKDILESSQRNNPANGITGLLCYSKPAFLQVLEGECEQVNETYHRIVQDERHHSPQIIECMPIRRRNFEVWSM

>Ycgf_Escherichia_coli_CAD6016340.1

MLTTLIYRSHIRDDEPVKKIEEMVSIANRRNMQSDVTGILLFNGSHFFQLLEGPEEQVKMIYRTICQDPRHYNIVELLCDYAPARRFGKAGM

>BlrB_Cereibacter_sphaeroides_ABA80440.1

ELVSLTYRSRVRLADPVADIVQIMRASRVRNLRLGITGILLYNGVHFVQTIEGPRSACDELFRLISADPRHQEILAFDLEPITARRFPDWSM

>TII0078_Thermosynechococcus_vestitus_BAC07631.1

LHRLIYLSCATDGLSYPDLRDIMAKSEVNNLRDGITGMLCYGNGMFLQTLEGDRQKVSETYARILKDPRHHSAEIVEFKAIEERTFINWSM

>OaPac_Oscillatoria_acuminata_pdb|4YUT

LTYISKFSRPLSGDEIEAIGRISSQKNQQANVTGVLLCLDGIFFQILEGEAEKIDRIYERILADERHTDILCLKSEVEVQERMFPDWSM

>BlrP_Escherichia_coli_P75990.1

MLTTLIYRSHIRDDEPVKKIEEMVSIANRRNMQSDVTGILLFNGSHFFQLLEGPEEQVKMIYRAICQDPRHYNIVELLCDYAPARRFGKAGM

>bPAC_Beggiatoa_sp_UPI000159043

MMKRLVYISKISGHLSLEEIQRIGKVSIKNNQRDNITGVLLYLQGLFFQILEGENEKVDKLYKKILVDDRHTNILCLKTEYDITDRMFPNWAMKT

>BlsA_Acinetobacter_V5VB82

RLCYASQRNEKNEDLLQDLRDILTEARDFNDLNGICGVLYYADNAFFQCLEGEQEVVERLFEKIQKDQRHYNIKWLCTYSIDEHSFQRWSMK

>Magnetococcus_WP_011712478.1

IFHLIYVSKTDVLKPGDLGDILFAARKNNPPLNLTGVLIYNSDFFFQLLEGPASNVEKMYKIICDDPRHIACKTLHTYTDYHRKFTQWSM

>Mycosarcoma_maydis_XP_011386128.1

LIQVAYTSSALTRHMPLDEVKSILRTSRINNTRQNITGLLLYRDGSFAQFLEGPTDAVDS

LYDTIERDDRHHGVVRVLRQSITKRDFRQWSMA

>Ustilago_trichophora_SPO19831.1

LIQVVYTSSARSRHMPIDEVKSILRASRLNNMGKGITGLLLYRDGSFAQFLEGPADAVDS

LYDRIERDPRHHGVIRVVRQKVTKRDFRQWSMA

>Kalmanozyma_brasiliensis_XP_016294803.1

LIQVVYTSSAKSRHMPLDEVKQILRASRSNNTALGITGLLLYRDGSFAQFLEGPADAVDS

LYDKIERDSRHHGVIRVLRQPVAKRDFRQWSMA

>Sporisorium_scitamineum_CDR98633.1

LIQVVYTSSARSRTMSLDEVKSILRASRINNTSKGITGLLLYRDGSFAQFLEGPADAVDA

LYDKIERDPRHHGVIRVLRQSVTKRDFREWTMA

>Moesziomyces_antarcticus_SPO43039.1

LIQTVYTSSARSRHMSTDEVKSILRASRTNNSRLGITGLLLYRDGTFAQFLEGPAEAVDA

LYDSIERDPRHHGVIRVLRQSVTKRDFKQWSMAF

>Pseudozyma_hubeiensis_XP_012190327.1

LIQVVYTSSARSRLMSIDEVKSILRASRANNTAKGITGLLLYRDGSFAQFLEGPAHAVDS

LYDTIERDQRHHGVIRVLRQSVTKRDFRQWSMA

>Moesziomyces_aphidis_ETS63445.1

LIQMVYTSSARSRHMSTDEVKSILRASRSNNSRLGITGLLLYRDGTFAQFLEGPADAVDA

LYDSIERDPRHHGVIRVLRQHVTKRDFKQWSMA

>Melanopsichium_pennsylvanicum_CDI51132.1

LIQVVYTSSARSRQMSNDEVKSILRASRFNNTGKGITGLLLYRDGSFAQFLEGPAHAVDS

LYDKIERDPRHHGVIRILRQSVAKRDFREWSMAF

>Sporisorium_reilianum_SJX60221.1

LIQVVYTSSARSRSLSLDEVKSILRASRLNNTSQGITGLLLYRDGSFAQFLEGPADAVDA

LYDKIERDPRHHGVIRVLRQSVTKRDFREWSMG

>Sporisorium_graminicola_XP_029742422.1

LIQVVYTSSASSRTLSVDEIKAILRASRLNNTSKGITGLLLYRDGSFAQFLEGPAYAVDA

LYDKIERDPRHRGVIRVLRQSVTKRDFREWSMA

>Ustilago_loliicola_KAJ1030449.1

LIQVVYTSSARSRHMTNDEVRSILRGSRANNTSKGITGLLLYRDGSFAQFLEGPVALVDE

VYDKIERDPRHYGVIRVLRQAVTKRDFRQWSMAF

>KAJ1030823.1_Ustilago_nuda

LIQVVYTSSARSRQMTKDEVKSILRTSRTNNTCKGITGLLLYHDGSFAQFLEGPAAAVDA

LYHKIEHDPRHHGVIRVLRQPVTKRDFKQWSMA

>Ustilago_hordei_XP_041409432.1

LIQVVYTSSARSRQMTKDEVKSILRTSRTNNTCKGITGLLLYHDGSFAQFLEGPAAAVDA

LYHKIEHDPRHHGVIRVLRQPVTKRDFKQWSMA

>Ustilago_bromivora_SAM71631.1

LIQVVYTSSARSRQMTKDEVKSILRTSRTNNTCKGITGLLLYHDGSFAQFLEGPAAAVDA

LYHKIEHDPRHHGVIRVLRQPVTKRDFKQWSMA

>Ustilago_tritici_KAJ1031905.1

LIQVVYTSSARSRQMTKDEVKSILRTSRTNNTCKGITGLLLYHDGSFAQFLEGPAAAVDA

LYYKIEHDPRHHGVIRVLRQPVTKRDFKQWSMA

>Testicularia_cyperi_PWZ02834.1

LLQVVYTSSAYSRHITITEIRSILQASRSNNESKDITGLLLYRDGSFAQFLEGPASAVDA

LYQKIERDPRHRGIIRVLRQPATKRDFSRWSMA

>Pseudozyma_flocculosa_SPO34781.1

LQVVYTSSALTRHVSREEISDILQHSRRNNARRGITGLLLYRDGSFVQFLEGPAHEVDGV

YQKIEADPRHRGIVRILRKTVEKRDFSKWEMA

>Sympodiomycopsis_kandeliae_CAO1613844.1

LTQIVYTSSASHRYLPSEELTSILHGSRKRNCENDITGLLLYRDGSFAQFLEGPNEAVQS

TFAKIETDQRHKGVVVVLNRSVKKRDFPTWRMG

>Microstroma_glucosiphilum_XP_025346216.1

LIQLVYTSSAKIRYLPRSELEGILASSRKTNARNDITGLLMYRDGSFAQFLEGPRSAVLD

TFARIRADRRHRGVIVVLKRPVEKRDFAEWRMG

>Ceraceosorus_guamensis_XP_025369106.1

LLQIVYASSSPERSPAAASVEQILRTSRRNNAAEDITGLLLFHDGSFVQFLEGPPANVKR

VYERIRKDDRHRGVIEILSVSVAK

>Tilletiopsis_washingtonensis_XP_025601205.1

LLHLVYTSSSPDRYMTRASIEQILATSRRNNSRVGVTGLLLYHDGSFVQFLEGPPAEVEA

VYQRICRDERHRGLIEIMRTRASQRSFREWSMAY

>Ceraceosorus_bombacis_CEH17329.1

LLQIVYASSSPERSPAAASVEQILRTSRRNNAAEDITGLLLFHDGSFVQFLEGPAANVKR

VYERIRKDDRHRGVIEIFSASVAKRSFADWSMA

>Jaminaea_rosea_XP_025361925.1

FDGTIGQDSLCQLVYTSSARDRYLTREQLQTLLNHSRHRNASKGITGLLLYRDGTFAQFL

EGPTHHVEALFETIRQDERHRGVIVVLKRQGLQKRDFDGWRMA

>Jaminaea_angkorensis_CAO1631867.1

LCQLVYTSSARDRYMARNELQTILSHSRKANASKGITGLLLFRDGTFAQFLEGPQQHVEA

LFERIQQDERHRGVIVVLKRQGVRRRDFAEWRMAY

>Savitreella_phatthalungensis_BFZ55319.1

LYQLVYLSSAEHLLEDSELVSILAASRRNNIKRNVTGLLLYHDGNFIQFLEGPAHQVDAL

YERISLDVRHRGVLRLLRREIAKRDFGEWTMG

>Parajaminaea_phylloscopi_CAO1635359.1

LYQLVYTSSSRNRYLTPDVLQSILVKSRSRNASNDITGLLLYRDGSFAQFLEGPRDAVCA

TFHRIEADERHRGVIVVLQREIERRDFDSWRMAF

>Phaffia_rhodozyma_CED85203.1

FQLVYCSSSISPAMPRDDLIDILSVCRRNNAAAAISGLLMYKDGQFVQFLEGPEHRVRRI

FTKINKDERHSGVYVLLEQKTNKRDFPQWSMA

>Taphrina_deformans_CCG84725.1

LYQLIYLSSSAGSFSKEDLTDILTTSRRNNTRQGITGLLLYHEGTIIQFLEGNESVIQNL

YNIIAMDTRHKGVLPLLKRKIERRDFGTWTMGF

>Fimicolochytrium_jonesii_XP_052927778.1

LRQIVYVSQARTDLTSRQLAAILNTARTNNDHNKISGMLVFNSGQFMQCIEGPWSEVEAL

MSGIKKDRRHEYMAILLDQAITDRDFDCWLMG

>Powellomyces_hirtus_KAI8913094.1

IRQLIYLSQAVTPSSAALAAILDSARRRNERDQISGILLYNNGQFMQCLEGNHDQVGAVY

NDIRKDDRHDHLAVLLDHTVSRRDFDCWLMGF

>Geranomyces_variabilis_KAJ3176981.1

IRQLVYLSQAVVSTPAALASILKSARSRNEREKISGMLLFNNGQFMQCLEGSPEKVGAVY

QAISKDDRHDHLAILLDHIIPRRDFDSWLMAF

>Geranomyces_michiganensis_KAJ3149842.1

IRQLVYLSQAVSSSPQDLASILKSARHRNERDQISGILLYNNGQFMQCLEGDPEKVGAVY

KDISRDKRHDHLAILLDHKIERRDFDAWLMA

>Thoreauomyce_shumboldtii_KAJ3009313.1

VRQLVYTSDAAPYLGPEDHLDILSSARSYNAVHDITGMLLHAASTFIQVLEGPSDEIEEV

YLRISRDPRHTRPVILLDALTHQRDFECWMMG

>Friedmanniomyces_endolithicus_KAK0331322.1

MKRLLYISTARAILPATELDELLLKSREANSRAGITGLLIVGGRRFLQVLEGADEAVCAT

YERIMRDPRHFALVKLHDKQVESRSFGTWDMRF

**Supplementary table**

| Name | Sequence |
| --- | --- |
| MmBLUF-T93Y-fwd | **ATCGAATCCGTTACCTATGCATCCTCTTTCACT** |
| MmBLUF-T93Y-rev | **AGTGAAAGAGGATGCATAGGTAACGGATTCGAT** |
| MmBLUF Fwd | **CGGGGATCCATGCTGCAACAAGCGTCT** |
| MmBLUF REV | **GTCCTCGAGTTACAGCGGGCGATCCATAAAG** |
| MmBLUF T-60 | **CAAGGATCCGCGAGCAGCCACAAC** |
| MmBLUF T-100 | **CGGGGATCCGCATCCACCAGCAACTAC** |

**Table S1: List of the primers used in the present study for site directed mutagenesis**

**Supplementary figure**

**Figure S1**

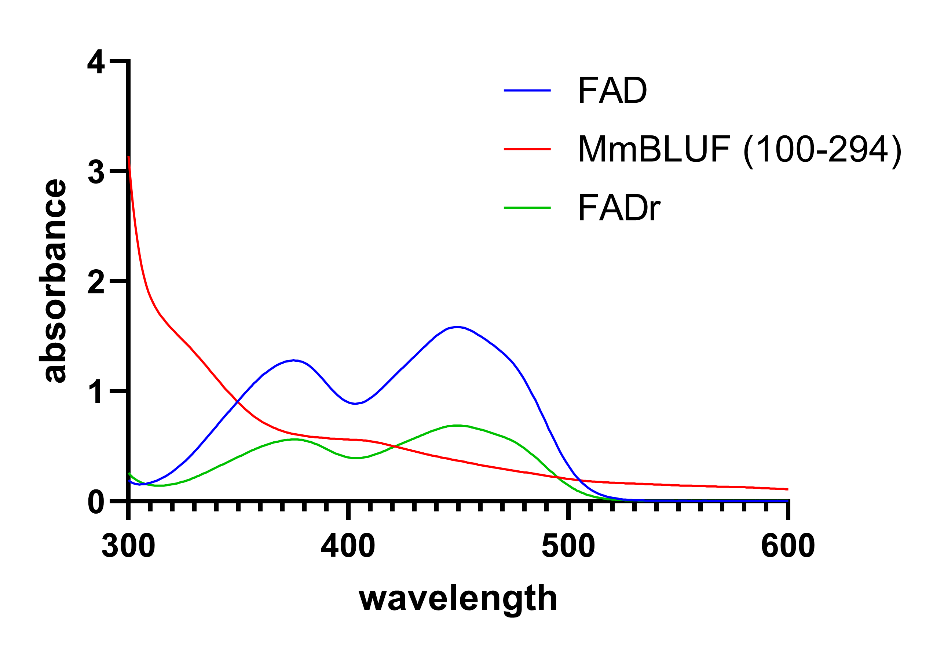

**Figure S1. UV-vis Spectra of free flavin di-nucleotide (FAD) overlayed with reduced FAD and MmBLUF (100-294).** Free flavin was reduced using sodium dithionite, which caused a decrease in its absorbance.

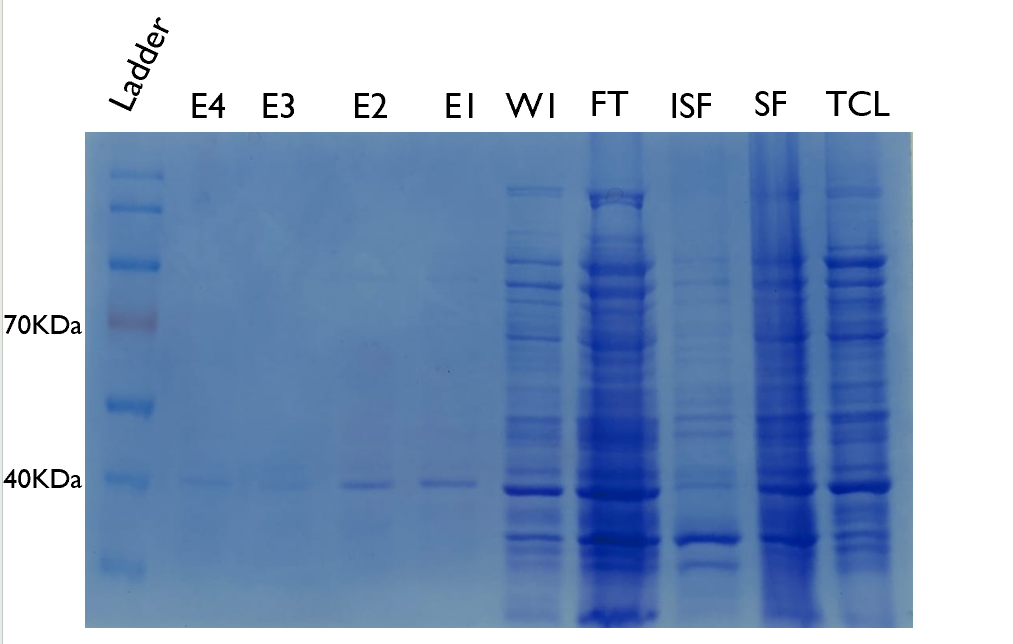
**Figure S2**

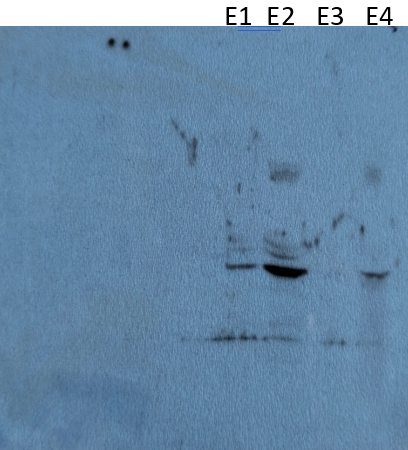

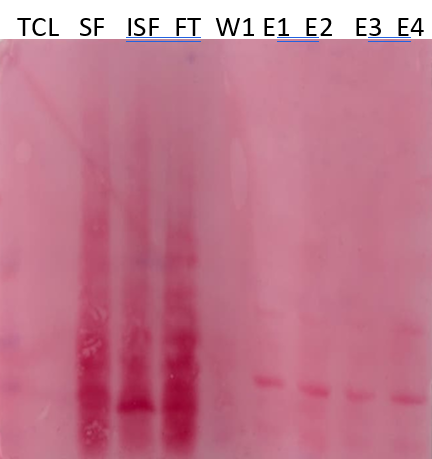

C

B

A

**Figure S2: SDS PAGE and immunoblotting of MmBLUF**

A: SDS PAGE TCL-Total cell lysate, SF-soluble fraction, ISF-Insoluble fraction, FT-flow through, W1-wash, E1-E4-Elutions.

B: Ponceau-stained gel showing Transfer of Proteins from Gel to Nitrocellulose membrane.

C: X-ray film, post western blotting
